# Polygenic Risk Scores Across Genomic Platforms for Reliable Breast Cancer Risk Stratification

**DOI:** 10.1101/2025.04.10.648141

**Authors:** Peh Joo Ho, Alexis Jiaying Khng, Joanna Hui Juan Tan, Pierre-Alexis Vincent Goy, Kayla Aisha Kamila, Zheng Li, Weang Kee Ho, Iain Bee Huat Tan, Dawn Qingqing Chong, Elaine Lo, Liuh Ling Goh, Hwee Lin Wee, Mikael Hartman, Rajkumar Dorajoo, Nicolas Bertin, Jingmei Li

## Abstract

**Purpose:** We evaluated differences in a 313-variant breast cancer polygenic risk score (PRS_313_) across genomic platforms and their impact on risk stratification.

**Methods:** We compared PRS_313_ derived from genotyping arrays (Global Screening Array [GSA], OncoArray-500K [OncoArray], Global Diversity Array [GDA], custom Axiom_PrecipV1 array [ThermoFisher]) and low-coverage genome sequencing (lc-WGS) in 2 cell lines and 92 individuals. Probes were designed for all variants on ThermoFisher (success rate: 259/313). Sanger sequencing was performed to profile indels. Concordance of high-risk classification (PRSscore>0.6) across platforms was assessed using Kappa statistics.

**Results:** PRS_313-lc-WGS_ was identical in the 4 cell line repeats. In saliva samples, indel concordance with Sanger sequencing varied widely (Kappa: 0.007–1.000). PRS313-ThermoFisher was predictable from other platforms using linear models, despite systematic differences. Greater agreement was observed between arrays with high imputation overlap (e.g., GDA∼GSA slope=0.986). Pre-calibration agreement in high-risk classification was moderate (Fleiss Kappa=0.552) and improved post-calibration (Kappa=0.650). Arrays with similar designs showed higher pre-calibration agreement (Kappa=0.745). Calibration narrowed high-risk proportions from 4–45% to 15–21% −28% were high-risk by any platform, while 8% were high-risk across all five.

**Conclusion:** Platform-specific biases affect PRS interpretation. Calibration enhances consistency in identifying high-risk individuals.

**STATEMENT OF SIGNIFICANCE:** This study compares the performance of a validated 313-variant breast cancer polygenic risk score across platforms, revealing systematic biases in risk stratification and raising concerns about including inconsistent indels in the model.

## INTRODUCTION

Interest in using polygenic risk scores (PRS) in clinical practice for better disease screening and diagnosis is steadily increasing.^1-3^ Although their validity is well-supported, the practical benefits of PRS in real-world settings remain insufficiently explored.^4-6^ In breast cancer risk prediction, PRS alone performs poorly for individual risk prediction and population stratification, indicating the need for integration with other risk factors.^7-11^ Another challenge is the variability in PRS distribution across populations, which may benefit from population-specific calibration.^12-15^

This study examines another source of variability: the impact of sequencing technology on PRS-derived breast cancer risk stratification. Sanger sequencing is recognized as the gold standard in many research and clinical applications due to its low error rate and high accuracy in detecting single nucleotide variants and indels.^16^ Arrays are key tools in PRS studies, with arrays offering cost-effective, high-accuracy genotyping for common variants.^17^ Recent advancements in low-coverage genome sequencing (lc-WGS) and improved imputation methods, such as GLIMPSE2, have improved rare variant imputation (MAF<1%), making lc-WGS a strong alternative to arrays. In addition, many countries are investing in efforts to sequence their populations.^1-3^ Previous studies have compared the overall performance and concordance of different genotyping and sequencing technologies; however, these analyses have generally averaged the results over the whole genome.^18,19^ It remains unclear whether these findings can be generalized to specific sets of variants.

Our study aims to fill this gap by focusing on a targeted 313-variant PRS (PRS_313_), the most widely studied breast cancer PRS.^20,21^ We assessed how different genomic platforms impact the accuracy and reliability of risk stratification for this specific set of variants. Genome-wide heritability estimates suggest that PRS loci account for approximately 40-45% of the heritability attributed to all common variants captured by genome-wide single-nucleotide variant (SNV) arrays.^20^ Discrimination statistics (AUC) for PRS_313_ within White European populations are approximately 0.63 to 0.653.^21,22^ PRS_313_ has been implemented in pilot risk-based breast cancer screening programs such as the Personalized Risk Assessment for Prevention and Early Detection of Breast Cancer: Integration and Implementation (PERSPECTIVE I&I) study^23^ and the BREAst cancer screening Tailored for HEr (BREATHE) study.^24^ To ensure reliable risk prediction and clinical application, we evaluate the degree of agreement between array and lc-WGS results, as the breast cancer PRS_313_ is primarily developed and validated on arrays.^25^

## METHODS

### Study population

Volunteers aged 21 to 80 years, who identified as Chinese, Malay, or Indian descent were eligible for our study. Between 12 December 2022 and 16 June 2023, 100 individuals signed up, of whom 94 completed informed consent. Two individuals withdrew before providing saliva samples. Our analytical cohort includes 92 individuals who provided ∼4 ml of saliva each (PAXgene Saliva Collectors from Qiagen, Part number: 769040, two tubes of 2 mL per individual), of whom 91 agreed to share their genetic data in the public domain.

We included two cell lines (GM18592, female; GM18609, male) to highlight potential differences due to poor DNA quality if any arose. Both cell lines were of cell type B-Lymphocyte, ethnicity Han Chinese, from the repository NHGRI Sample Repository for Human Genetic Research.

This study was approved by the A*STAR ethics board (reference number: 2022-062, approval date: 14 September 2022).

### DNA extraction

Saliva samples were pre-heated at 50°C in an air-incubator overnight before the purification of genomic DNA using prepIT•L2P reagent (Part number: PT-L2P-45) from DNA Genotek, according to the manufacturer’s recommendations.

### “Ground truth”: Direct genotyping and Sanger sequencing

We designed primers for all 313 variants in the breast cancer PRS panel using ThermoFisher Scientific’s Applied Biosystems™ Axiom™ arrays (Axiom_PrecipV1) (**Supplementary Table 1**). However, 54 primers (17%) failed quality checks. In total, 259 variants (235/265 SNVs and 24/48 indels) were successfully genotyped on ThermoFisher (**Supplementary Figure 1**). While primers for all 48 indels in the PRS panel were designed, four indels could not be successfully designed for Sanger sequencing validation. The remaining 44 indels were validated through Sanger sequencing (**Supplementary Table 1**).

Due to the large number of experiments required per individual and the limited amount of saliva samples available, we prioritized genotyping and low-coverage lc-WGS experiments over Sanger sequencing validation. As a result, Sanger sequencing was performed for only 87-89 individuals (3 had no DNA for Sanger sequencing; 2 had insufficient DNA for all indels).

### Genotyping and imputation on commercially available arrays

In addition to the custom ThermoFisher array, genotyping was done on three Illumina arrays: 1) Infinium Global Screening Array v3.0 (GSA), 2) Infinium OncoArray-500K v1.0 (Rev. C) (OncoArray), and 3) Infinium Global Diversity Array v1.0 (GDA) **(Additional Materials Section 1.1)**. Imputation using Minimac4 v1.7.4 was done on the Michigan Imputation Server (https://imputationserver.sph.umich.edu/index.html#!, accessed in February 2024).^26,27^ The imputation settings used are detailed in **Supplementary Table 2 (Additional Materials Section 1.2)**.

#### Low-coverage genome sequencing (lc-WGS)

Samples from participants were sequenced in 4 lanes, with 23 samples per lane. HapMap cell lines (GM18592 and GM18609) were included in each lane as controls. The raw sequencing data generated from lc-WGS were processed using the DRAGEN germline workflow (v3.7.8). Sequencing reads were aligned to the GRCh38 reference genome. Imputation of lc-WGS data was done using the 1000 Genome Projects Phase 3 reference panel (same as for the arrays, downloaded from^28^ http://ftp.1000genomes.ebi.ac.uk/vol1/ftp/data_collections/1000G_2504_high_coverage/working/20201028_3202_phased/) We created the binary reference panel using Glimpse2_chunk and Glimpse2_split_reference command. GLIMPSE2 was then used for the imputation of the lc-WGS dataset using this binary reference panel.

### Polygenic risk score (PRS) calculation

PRS_313_ was calculated as the weighted sum of effect alleles from 313 breast cancer variants, as previously described (**Supplementary Table 3**).^20^ This was performed with PLINK 1.9 using the “scoresum” option.^29^

### Statistical analysis

#### Assessment of imputation quality and consistency of PRS_313_ across platforms using cell lines

To assess the impact of imputation on the consistency of PRS_313_ calculations, we compared PRS_313_ across platforms. Cell lines were used for this purpose to minimize the potential influence of DNA quality. For array-based genotyping, PRS_313_ was calculated using various subsets of variants at different imputation quality thresholds (from rsq≥0 to >0.9, at 0.1 intervals). Lc-WGS used a different method for imputation (GLIMPSE2) where changing the imputation quality will not change the subset of variants consistently across all individuals.

#### Assessing agreement, correlation, linear predictions, and high-risk classification in saliva samples

Fleiss’s Kappa statistic was used to assess the agreement in variant calling between the different platforms. For indels, Kappa was calculated between each platform and the results from Sanger sequencing.

We assessed the correlations and linear predictions of PRS_313_ calculated from different platforms to evaluate the need for calibration. Two approaches were used: 1) no variant exclusion (i.e. PRS_313_ calculated with variants with rsq≥0), and 2) only high-quality imputed variants (rsq>0.9).^18^ Repeated calculations were also performed for rsq>0.3 and rsq>0.8, which are other commonly used thresholds for imputation variant quality control.^18^

In our hypothetical scenario, not representative of the general population, we compared high-risk classification differences between PRS_313_ derived from the different platforms. The proportion of high-risk individuals identified depends on the mean and standard deviation (SD) of the PRS_313_.^22^ The threshold for high risk was determined based on the PRS_313_ distribution in non-breast cancer controls, which followed a Gaussian distribution with a mean 0.130 and a standard deviation of 0.565).^8^ The 80th percentile of the PRS_313_ distribution in these controls is ∼0.6. Hence, in our analyses, individuals with PRS_313_ (scoresum) greater than 0.6 were classified as high-risk. Calibration involved adjusting the mean of the PRS_313_ distribution to match that of the reference population’s mean of 0.130. Fleiss’s Kappa statistic was used to study the agreement in high-risk individuals identified across the platforms.

#### Evaluation of additional disease risk PRS across platforms

To test generalizability of findings to other PRS, we assessed PRS for other diseases with different numbers of variants included, such as prostate cancer (PGS000662, 269 variants)^30^, colorectal cancer (female [PGS000055, 76 variants^31^], male [PGS000734, 95 variants^32^]), hypercholesterolemia (PGS000889, 9,008 variants)^33^, and coronary heart disease (PGS003446, 537,871 variants)^34^. The corresponding PRS score files were obtained from the PGS catalog (accessed in May 2024).^35,36^

The GRCh38 reference genome was used for all analyses. Statistical analysis was done using R version 4.2.2.

## RESULTS

### Characteristics of 92 individuals

A total of 92 participants donated saliva samples. The median age was 32 years (interquartile range [IQR] of 26 to 38 years), 71% (n=65) were females, 68% (n=63) identified themselves as Chinese, 22% (n=20) identified as Indian and 10% (n=9) identified as Malay (**Supplementary Table 4**).

### Directly typed variants

Among 2,734,610 unique variants typed across the different arrays, 28,646 (1%) were common across all four (**Supplementary Figure 2**). Less than 50% of the PRS_313_ variants were present on any of the Infinium arrays: 64 (20%) on GSA, 99 (32%) on OncoArray, and 82 (26%) on GDA (**Supplementary Figure 2, Supplementary Table 3**). All variants found were SNVs. Among the 313 variants, 259 (83%) were successfully designed on ThermoFisher, including 235 SNVs and 24 indels.

### PRS313 using cell lines

Figure 1. shows variations in calculated PRS (y-axis) for 1 to 4 repeats per cell line. Four technical repeats per cell line were performed for lc-WGS (Repeats 1 to 4). Genotyping chips have a fixed capacity. After assigning slots to the 92 unique samples, the remaining slots allowed for two technical replicates (Repeats 1-2) per cell line on GSA, OncoArray, and GDA. For ThermoFisher, the fixed capacity allowed for the inclusion of only one sample per cell line.

**Figure 1.**
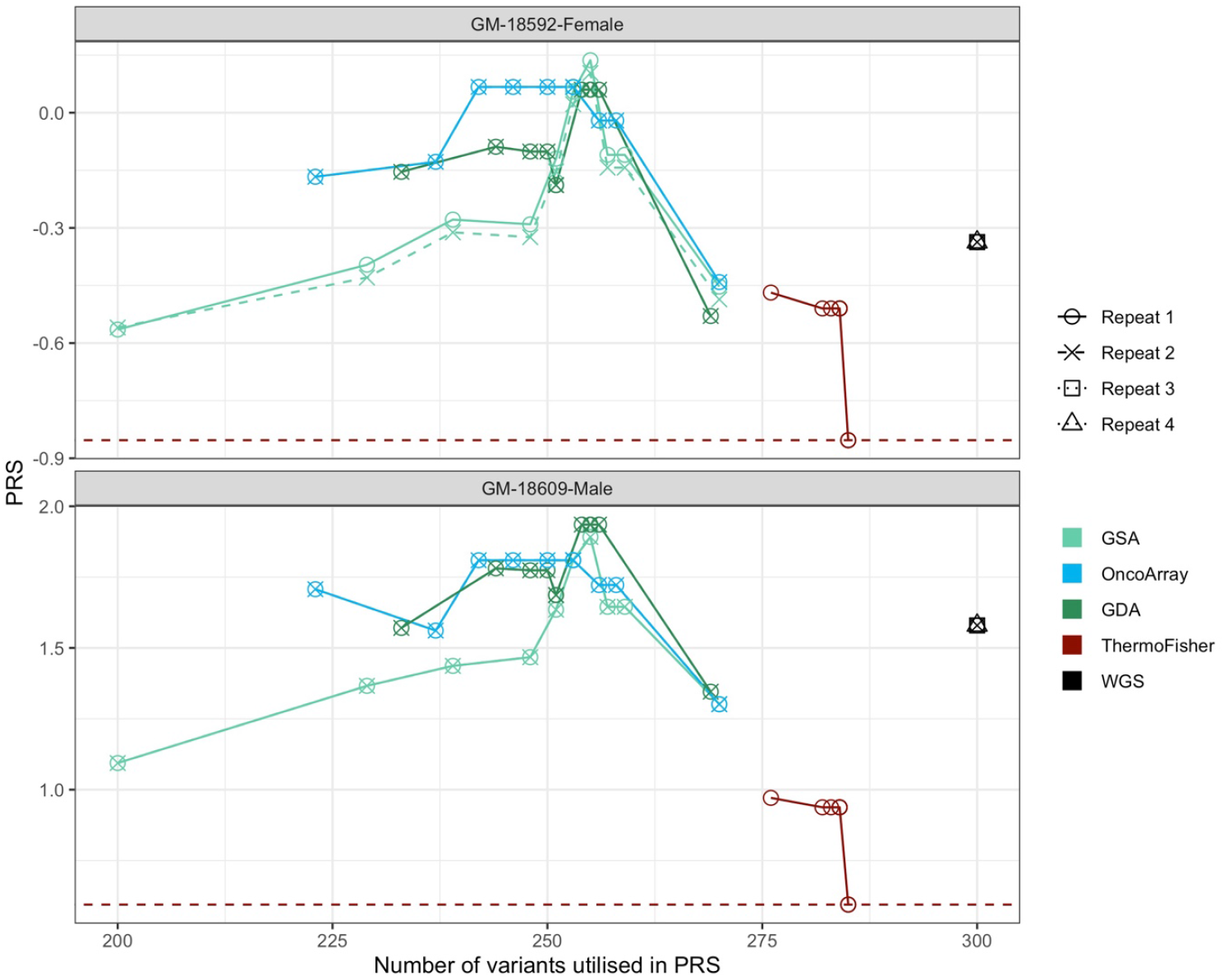
PRS313 derived from cell lines, 1-4 repeats depending on platform used (indicated by the symbols). In black is the result from the repeats 1-4 (represented by 4 different symbols, one in each sequencing lane) for low-coverage whole genome sequencing (lc-WGS). Imputation quality is not measured per variant, all 300 variants (13 could not be imputed) were used for the PRS calculation. Two repeats were performed for each cell line (together with the 92 samples on each array) for GSA (turquoise), OncoArray (blue) and GDA (green). In red, is the result of repeat 1 (no additional repeats possible due to allocation on array) from ThermoFisher, where 256 variants were directly genotyped. For arrays, we calculated PRS313 with different imputation quality thresholds (rsq 0 to 0.9 at 0.1 intervals). The number of variants utilized in PRS (x-axis) decreases with higher imputation quality (i.e. higher imputation [left] to lower imputation quality [right]).

Post-imputation, between 86% and 96% of the 313 variants were available for analysis (regardless of imputation quality). Specifically, 269 (86%) variants were available for GDA, 270 (86%) for GSA and OncoArray, 285 (91%) for ThermoFisher, and 300 (96%) for lc-WGS (**Supplementary Figure 1**). lc-WGS yielded identical PRS_313_ values (to four decimal places) across four repeats. ThermoFisher’s PRS_313_, primarily based on directly genotyped variants (91%), was systematically lower than that from Infinium arrays or lc-WGS when using the same imputation quality threshold. PRS_313_ results from Infinium arrays were similar when all variants were included (rsq≥0) but diverged when restricted to high imputation quality variants (rsq>0.6), likely due to differences in variant subsets (**Figure 1**).

**Supplementary Figure 1** shows the number of genotyped/ imputed SNVs/ indels at selected imputation quality levels (rsq≥0, >0.3, >0.8, or >0.9).

### Sanger sequencing for indels

Sanger sequencing was performed for 87-89 individuals (**Supplementary Table 5, Figure 2**). The four indels not validated by Sanger sequencing were on (chr:position:A1:A2) chr1:168201814:C:CA, chr4:125831837:AAT:A, chr16:3958541:C:CAAAAA, and chr19:19406245:CGGGCG:C (Supplementary Table 1). However, chr1:168201814:C:CA was typed on ThermoFisher. chr1:168201814:C:CAAAAA and chr19:19406245:CGGGCG:C were available in the post-imputation lc-WGS data. We did not observe variations in 17/44 indels that was Sanger sequenced, hence pairwise agreement with other platforms cannot be calculated (NA).

**Figure 2.**
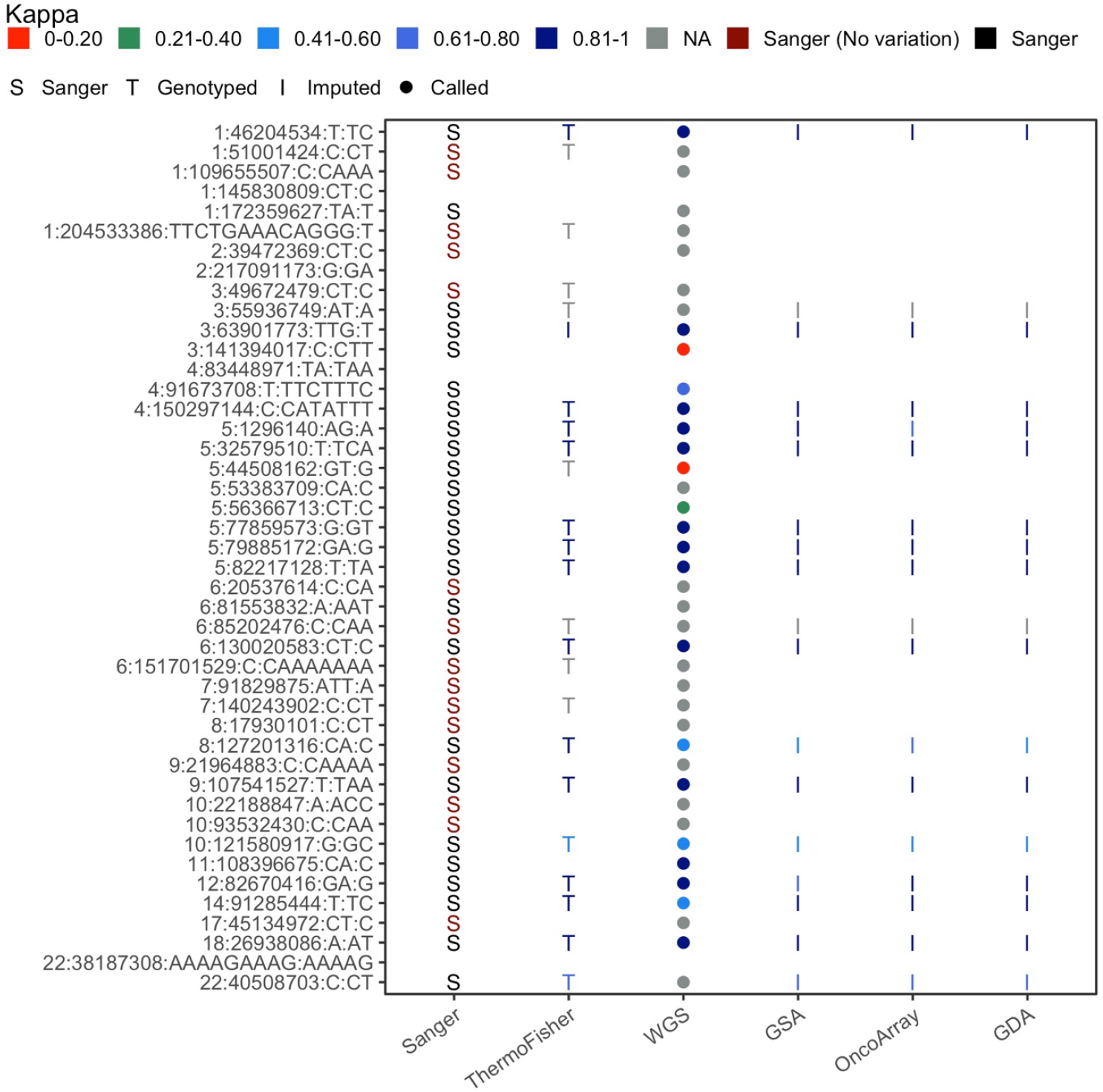
List of indels (44 of 48) profiled using Sanger sequencing. Pairwise agreement (Kappa) of indels between Sanger and platforms (ThermoFisher, lc-WGS, GSA, OncoArray and GDA) are presented.

No indels were genotyped on the Infinium arrays but 18 indels (41% of the 44 indels) were imputed. Agreement between Sanger sequencing and imputed Infinium variants ranged between 0.515 (chr8:127201316:C:CA on GSA) to 1 (**Supplementary Table 5**). Of the 30 indels imputed in lc-WGS (MAF>1%), 7 indels had Kappa<0.5 and Kappa was not calculated for 8 indels (i.e. no variation in Sanger) (**Supplementary Table 5**).

### Concordance of variants across platforms

Among the PRS_313_ variants in saliva samples, 266 (85%) were genotyped or imputed across all platforms (arrays and lc-WGS). Almost perfect agreement (Kappa>0.8)^37^ between all platforms was observed for 78% (n=206) of these 266 variants (**Supplementary Table 6**). However, 15 SNV showed slight to no agreement (Kappa≤0.2). The median concordance (Fleiss Kappa) between all platforms was 0.962 [IQR: 0.860 to 0.993]. Between arrays, it was 0.972 [IQR: 0.846 to 1]. Excluding ThermoFisher, median concordance between arrays increased to 0.984 [IQR: 0.885 to 1] (**Supplementary Table 6**).

### High PRS_313_ correlation across platforms, but different risk scores are obtained for the same individuals

Pairwise correlation between PRS_313_ derived from different platforms (rsq≥0) ranged from 0.754 (lc-WGS∼GSA) to 0.940 (GDA∼OncoArray) (**Figure 3, Supplementary Table 7**). PRS_313_ calculated from ThermoFisher (91% directly typed variants) could be predicted using PRS_313_ from the Infinium arrays and lc-WGS via linear models. However, PRS_313_ with a high proportion of imputed variants (Infinium arrays and lc-WGS) tended to overestimate PRS313. The GDA∼GSA pair exhibited a slight shift (intercept=-0.013) in mean but maintained a slope close to 1. (**Figure 3**). When restricting the PRS calculation to variants with higher imputation quality (rsq>0.3, >0.8, or >0.9), similar linear relationships were observed (**Supplementary Figure 3**).

**Figure 3.**
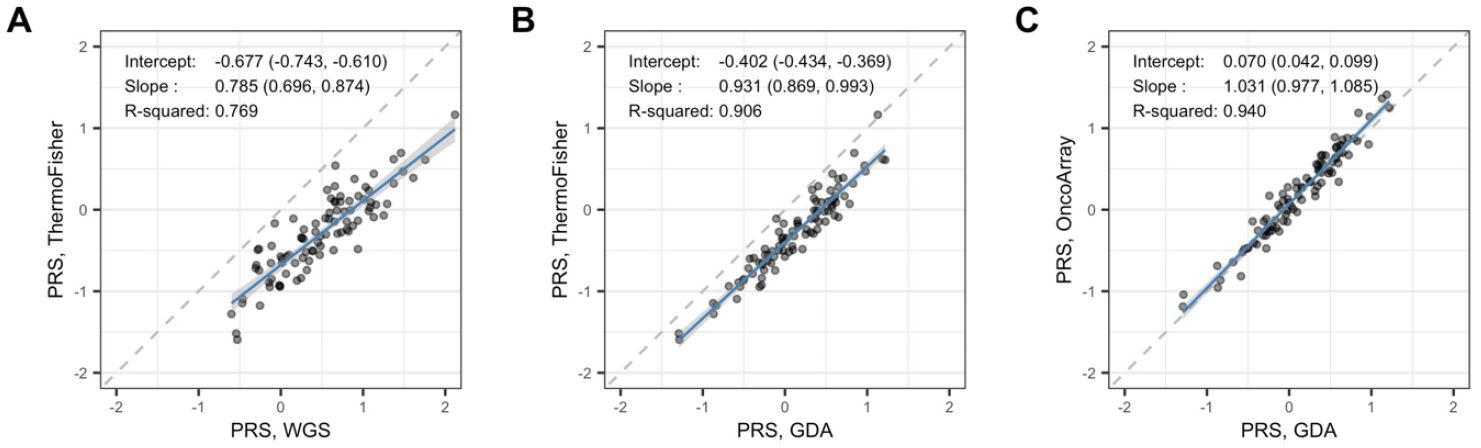
Selected pairwise linear association of PRS313, computed using all variants that are genotyped or have imputation quality rsq≥0. A) Using lc-WGS to predict ThermoFisher (ground truth). B) GDA (185s/313 variants imputed) prediction of the ground truth (ThermoFisher). C) GDA PRS (185/313 variants imputed) prediction of OncoArray PRS (171/313 variants imputed). The intercept and slope with respective 95% confidence intervals (denoted by blue line), and correlation (r-square) are reported. All pairwise comparisons are presented in **Supplementary Figure 3 and Supplementary Table 7**.

PRS_313_ variability (standard deviation [SD]) across all platforms and imputation quality thresholds ranged from 0.464 to 0.565 (**Figure 4, Supplementary Table 8**). Without calibration, at rsq≥0, 42 (46%) were high-risk by any platform, with 4 (4%) identified as high-risk by all five platforms. Without calibration substantial agreement (Fleiss Kappa 0.61 to 0.80) among arrays and lc-WGS was observed when the classification of high-risk was performed without the restriction of imputation quality (rsq≥0). Moderate agreement (Fleiss Kappa 0.41 to 0.60) was observed when imputation quality was implemented (lowest *K*_rsq>0.9, infinium_=0.381 to highest *K*_rsq>0.8, infinium_=0.559) (**Supplementary Table 9**).

**Figure 4.**
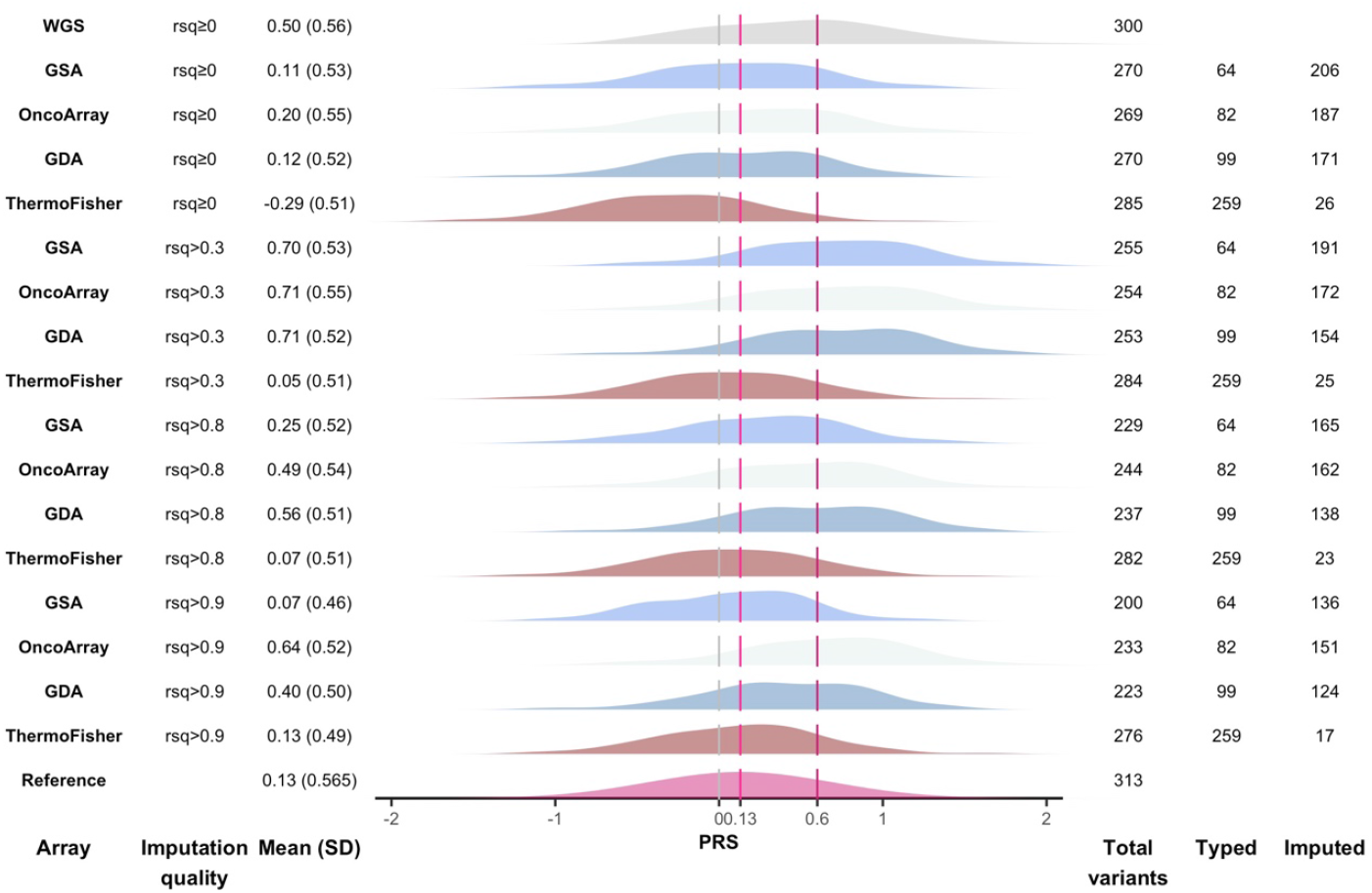
Distribution of breast cancer polygenic risk scores (build GRCh38). Imputed variants must be above the R-square (Rsq) threshold.

### The chosen platform affects who is identified as high risk, especially those not at the extremes of the risk spectrum

Post-calibration, a total of 26 (28%) unique individuals were high-risk by any platform, with 7 (8%) identified as high-risk by all five platforms (**Figure 5**) The high-risk proportion ranged between 12% and 21% across platforms and imputation quality, closer to the expected 20% based on the threshold selected from the reference population (**Supplementary Table 8**). Agreement improved (lowest *K*_rsq>0.9, infinium_=0.599 to highest *K*_rsq>0.8, infinium_=0.808) after calibration (centering the mean to that of the reference population of 0.130) (**Supplementary Table 9**).

**Figure 5.**
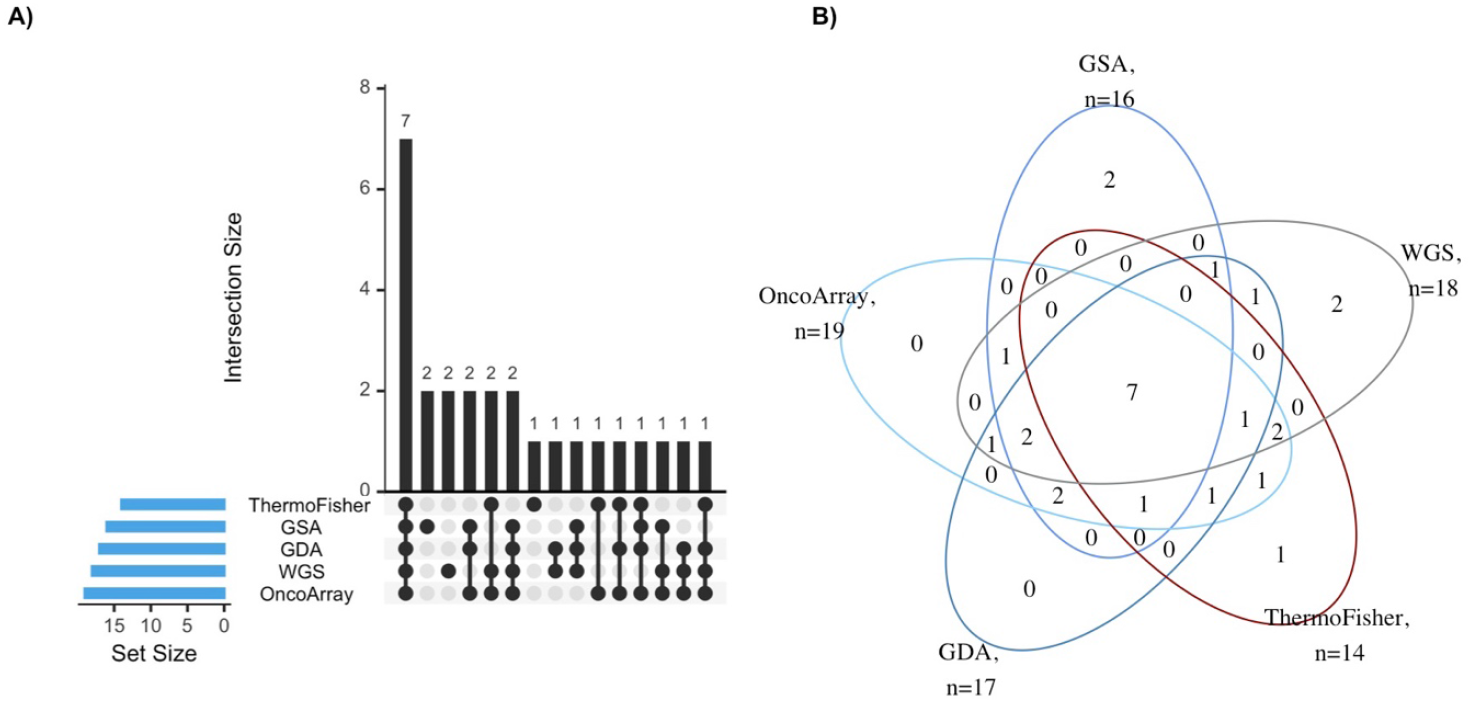
High-risk individuals (n=26 of 92) identified by the platforms, at rsq≥0. A) A matrix layout for intersections that has at least one individual. Set size denotes number of high-risk individuals identified on each array. Dark circles indicate sets that are part of the intersection. B) The same information on the overlaps of high-risk individuals identified, depicted in a Venn diagram.

### Repeating the analysis with variants common across all platforms (Additional Material Sections 2 and 3)

Even with the same set of variants used, PRS derived from Thermofisher (PRS_313_ with predominantly genotyped variants) differed from PRS_313_ derived from Infinium arrays (PRS_313_ with predominantly imputed variants). However, the correlation between PRS_313_ derived from Infinium arrays rose to 0.901 (GSA∼OncoArray), 0.933 (GDA∼GSA), and 0.945 (GDA∼OncoArray). Additionally, using variants common across all arrays resulted in higher agreement across all platforms in identifying high-risk individuals; of the 24 high-risk individuals identified by at least one platform, 11 (46%) individuals were identified by all five platforms.

### PRS calculated from a subset of SNVs

Agreement of indels across platforms is poor (**Figure 2**). We thus excluded indels and used 251 (GDA), 252 (GSA and OncoArray), 260 (ThermoFisher), and 256 (lc-WGS) SNVs (rsq≥0) for PRS_313_ calculation (**Supplementary Table 8**). The standard deviation (range: 0.472 and 0.509) was smaller when compared to 0.506 to 0.565 when indels were included. The smaller SD resulted in fewer individuals identified as high-risk (13-18%, compared to 15-21% previously) (**Supplementary Table 8**). Correspondingly, agreement in the high-risk individuals identified by the different arrays was improved (Kappa_all_arrays_=0.708, previously 0.668) (**Supplementary Table 9**). In contrast, among the Infinium arrays, the agreement in the high-risk individuals decreased (Kappa_Infinium_=0.697, previously 0.739; at rsq≥0) (**Supplementary Table 9**).

### Generalizability of findings to other PRS

The proportion of variants genotyped on commercial arrays is low for PRS scores that include a large number of variants **(Supplementary Table 10)**. Post-imputation among the five PRS, the proportion of available variants ranged from 84% to 97% for GSA, 89% to 97% for OncoArray, 89% to 97% for GDA, and 89% to 97% for ThermoFisher. lc-WGS with imputation by GLIMPSE2 had the largest range, from 94% to 99% **(Supplementary Table 10)**. platforms with the largest difference in the proportion of variants available saw large differences in mean PRS calculated. In particular, PGS003446 (coronary heart disease) with a large number of variants (537871 variants) was most affected; the highest proportion of available variants was for lc-WGS [97%] and the smallest for GSA [88%]. However, the correlation between coronary heart disease’s PRS calculated remains high (r-squared range from 0.926 to 0.968) (**Supplementary Figure 4**). PRS with fewer variants were more likely to have a lower correlation between arrays (**Supplementary Table 11**).

## DISCUSSION

In an ideal setting, PRS_313_ is calculated using directly genotyped variants, as this would provide the most accurate and reliable measurement for making personalized healthcare decisions. However, due to practical constraints such as cost and logistics, selective genotyping and imputation of missing genotypes are often performed.^38,39^ While these methods help increase the number of available variants, they also introduce variability in the proportion of directly genotyped variants and imputation success rates. Our study highlights key issues and opportunities in the comparability and utility of PRS_313_ across different genotyping arrays and sequencing platforms. We observed variability in concordance between genotyped indels and Sanger sequencing. Although PRS_313_ values from different platforms are highly correlated, varying risk scores were obtained for the same individuals. Calibration improved risk classification agreement, reducing variability in the proportion of high-risk individuals identified across platforms (from 4-45% pre-calibration to 15-21% post-calibration), aligning more closely with the expected 20% from a reference population. However, high-risk individuals identified across different platforms remained largely inconsistent.

We first examined PRS_313_ generated from cell lines, which experience fewer issues with variability in DNA quality. The availability of PRS313 variants post-imputation varied across platforms, with lc-WGS showing the highest proportion (96%) of available variants, followed by ThermoFisher (91%), GDA, GSA, and OncoArray (86%). Besides high variant availability, lc-WGS demonstrated excellent reproducibility, producing identical PRS_313_ values across four technical repeats. It is important to note that the discrepancy in variant availability will affect PRS_313_ calculation, particularly if variants with higher weightage for risk prediction are unavailable or of lower imputation quality on certain platforms.

When examining PRS_313_ in 92 real-world biological samples, a key issue we revealed is the inadequate capture of PRS_313_ indels using commercial off-the-shelf arrays. Our study found that none of the commercially available arrays included any of the 48 indels from PRS_313_. Only half of the indels (24/48) passed probe design on the custom ThermoFisher array, and 44 primers were successfully designed for Sanger sequencing. Even between the directly genotyped indels, there was considerable variability in agreement. On the other hand, the high agreement observed with Infinium arrays (where all indels were imputed) may reflect the influence of the common imputation reference panel, rather than the inherent accuracy of array-based methods.^40,41^ These difficulties highlight the importance of carefully considering the inclusion of indels, especially when PRS_313_ includes a substantial proportion of indels, as this may significantly affect the accuracy of breast cancer risk prediction.

The variability in PRS_313_ across different platforms has clinical implications for risk stratification. Although there is a strong correlation between platforms, the risk scores can differ, especially for individuals not at the extremes of the risk spectrum.^25^ Agreement is better without restricting imputation quality, presumably because more data points are included. However, inaccuracies may result due to less reliable variants. In contrast, restricting to high-quality variants may improve prediction accuracy but result in fewer variants, thus reducing overall agreement in risk classification. While calibration improves consistency and reduces discrepancies by aligning risk classifications closer to the reference population, the results still indicate that 28% of individuals were identified as high-risk by at least one platform, and only 8% were identified by all five platforms. This suggests that while calibration helps, platform-specific PRS_313_ variability remains a challenge. To the layperson, this result matters because it shows that the test used to estimate a woman’s breast cancer risk can give different results depending on which method is used.

This study is the first to examine the consistency of PRS_313_ called across platforms, especially indels that were previously mostly imputed and not genotyped. The combination of Sanger sequencing, three commercial Illumina Infinium arrays, one custom ThermoFisher array, and lc-WGS provides a diverse and comprehensive selection of technologies for genotyping variants for PRS calculation. Efforts were made to design all PRS_313_ variants on the ThermoFisher array; Sanger sequencing was performed to validate all indels. A limitation of this study is that the imputation panel used was based on the 1000 Genomes Project, which may have limited sensitivity for less common variants in our Asian population. Nonetheless, this panel was applied consistently across all arrays and lc-WGS.

In this study, we found that the risk profiles based on the most validated breast cancer PRS can vary depending on the method. Women who are borderline high-risk are particularly affected. This means a woman’s risk could be classified differently depending on the platform used to generate the genetic data. Although we can adjust the results to make them more consistent, some differences still remain, which can impact decisions about screening and preventive care. In practice, it is important to realize that the method used can make a difference. It may not be feasible to derive PRS from multiple platforms or the same platform as the reference population, however, obtaining the PRS distribution of the first batch for calibration should be done before risk classification.

## ACKNOWLEDGEMENT

This work was supported by the A*STAR Computational Resource Centre through the use of its high-performance computing facilities.

